# A Study of Secondary School Students’ Participation in a Novel Course on Genomic Principles and Practices

**DOI:** 10.1101/2020.05.22.110361

**Authors:** Adam Stefanile

## Abstract

This paper presents the design, development, and validation of a study among secondary school students’ participation in a novel course on genomic principles and practices by analyzing and documenting evidence of their participation, and educational outcomes, in a novel course on genomic principles and practices. A mixed methods approach, using qualitative and quantitative methods, was used to address three research questions. 1) Based on affective evidence, how did secondary school students perceive and critically judge, content topics learned in a course on modern genomic principles and practices? 2) Based on cognitive evidence, how much of the content did secondary school students learn when they participated in a course on modern genomic principles and practices? 3) Using individual interview evidence, what are the major perceptions that the secondary school students expressed throughout the duration of the course? The participants were provided an opportunity to comment on the course through individual and collaborative interviews, in order to find out to what extent they perceived the course to be interesting and challenging. Future inquiry expanding from this research would help to establish the foundational pathway for designing a more inclusive genomics curriculum. In conclusion, the course offered real-life/real-world applications that encourage all students to conceptualize genomics, human health, diseases, medicine, ethics, beliefs, research, and careers.

## Introduction

Genomics is without a doubt one of the most groundbreaking scientific advances of the millennium, and scientists and science educators have endeavored to increase genomic education and awareness by highlighting a myriad of advancements in genomics, human health, and STEM [1–3]. Moreover, genomic education, has the potential to impact students and society exponentially. While concern about the state of science education and STEM education in America has been well documented [4], genomic education in the U.S. is unfortunately inadequate, in both quantity and quality [5–7].

The emergence of genomic research in the twentieth century has had a long and storied history of visionary research ideals in controversy on its merits and implications. In the early 1980s Renato Dulbecco was one of the first pioneers who researched the Human Genome Project (HGP) and supported the sequencing of human genomes to better understand cancer [8]. During that time (mid 1980s) the majority of biologists did not support the HGP, declaring that it was “bad science” and it was an arduous task that should be researched independently as opposed to collaboratively [8]. Many scientists also believed that the resources allocated towards the HGP would not justify the means [8–10]. One of the main objectives of the HGP has been, and continues to be, the generation of large, publicly available, comprehensive sets of reagents and data (scientific resources or ‘infrastructure’) that, along with other new, powerful technologies, comprise a toolkit for genomics-based research [11]. In fact, governmental agencies are consistently interconnecting and expanding their research in order to implement genomics into the public eye and mainstream media. These agencies include, but are not limited to, the following: Center for disease Control (CDC), Department of Energy’s (DOE), National Human Genome Research Institute (NHGRI), National Center for Biotechnology Information (NCBI) and Food and Drug Administration (FDA); and educational institutions. The future of genomics rests in the hands of our students; and the need to further establish genomics in high school curricula for the purpose of educating our future scientists is boundless [11–12].

Genomics is generally referred to as the study of the content, structure, organization, and functioning of an organisms’ genome. It is also concerned with the material contained in the genome that composes an organism and the analysis of multiple genes that interact with each other. However, genetics refers to the study of a singular gene of an organism. For over twenty years genomics has permeated many fields of science, and being that it is a relatively new and emerging field, it is an ideal discipline for designing and implementing it into an existing or new science curriculum. Genomics, specifically, is a progressively emerging field throughout almost every domain in science. The development of innovative technologies and analytic tools associated with genomics has improved many aspects of human condition and has changed the way that science research is being conducted today. This includes, but is not limited to, medicine, inheritance, pharmacology, diseases, forensics, reproduction, agriculture, and evolution.

These areas of research have, in turn, influenced ethical and social concerns, and stimulated the development of privacy laws and policies. There is overwhelming research supporting genomics education and how it is progressively expanding in the 21^st^ century from a hands-on laboratory-based science (e.g., micropipetting and sample preparation) to a cutting-edge computerized database (e.g., GENBANK, BLAST (Basic Local Alignment Search Tool), Medline) [13–15].

Over the last twenty years it has become increasingly common to read and watch news that is associated with genomic topics; and this requires society to be updated with scientific understandings and familiarity with modern principles, practices, and terminology in the field [16]. However, it should be noted that quite often, popular entertainment and mainstream media quite often mendaciously perpetuate incorrect genetic and genomic terminology to misinform the general public [17].

While acknowledging these advancements and benefits that genomics has on society [11, 15] there is very little attention to implementing genomics at the secondary school level [14, 37] level despite the overwhelming educational and career benefits. One of the current problems across the nation as it relates to science education is the lack of implementation and integration of modern biological principles, specifically genomics, into the secondary school curriculum. Secondary school students will be both the users and researchers of genomic information for the future, providing that secondary school educators have information and materials about genomics and its implications for society, to use in their classrooms [11].

There have been national efforts from organizations such as the National Institute of Health (NIH), the American Society of Human Genetics (ASHG), NHGRI, Howard Hughes Medical Institute (HHMI), the Jackson Laboratory (JAX), and Cold Spring Harbor Labs/DNA Learning Center (CSHL/DNALC) to improve genomic and science education and genomic and scientific literacy at the secondary school level. The level of science literacy of students in the United States, for instance, has been a source of concern to policymakers, educators, and citizens over the past decades, resulting in repeated calls and proposed strategies for raising students’ science proficiency [16]. Moreover, the physical resources are readily available and affordable for teachers and schools to adopt, and promote educationally, for student access, exposure, and learning. Modern biological and genomic topics such as: pharmacogenomics, genetically modified organisms (GMOs), Genome-Wide Association Studies (GWAS), epigenomics, genome sequencing, and genome editing are easily accessible via genomic educational resources and interactive websites (S1Fig).

Historically, genetics has been minimally taught at the middle school level, or taught as part of a biological unit at the secondary school level [18] and more typically taught in depth and as an elective course at the college level. The biology units usually encompass cells, chromosomes, genes, and alleles, dominant and recessive factors, genetic probability, independent assortment, gene segregation, and monogenetic diseases. Currently, on a smaller scale, some universities and institutions have developed a genomics curriculum and has been implemented to doctoral students, biology majors, pre-and-post-medical students, physicians, nurses, and pharmacists [19–20]. The advancement in many aspects of genomic education has personalized, and continues to personalize, genomic principles and practices to provide a practical learning opportunity for students to not just discuss genomics in abstract, but to engage in a more active and relevant way that makes it personal for them [21].

These findings provide evidence involving an educational gap that currently exists in genomic education at the secondary school level, which, has been identified and acknowledged by personal interviews conducted with numerous scientists at the American Museum of Natural History (AMNH), the Jackson Laboratory, and NIH; and science educators at ASHG and Teachers College, Columbia University. Acquiring adequate genomic science education in K-12 is needed as a means to increase long-term genomic health literacy in our society [17].

Successful dissemination, understanding, adoption and adherence to genomic health recommendations will require an elevation of the genomic literacy of the public in the context of public health genomics– to promote the appropriate translation of the new science of genomics into health benefits to individuals and populations, and for evaluating the impact of genomic information on health care and disease prevention [17].

The purpose of this research was to document and analyze evidence of secondary school students’ participation and educational outcomes in a novel STEM course on genomic principles and practices: Introduction to Genomics. One of the key guiding assumptions is by providing access and exposure to learning genomics the participants’ level of content knowledge will increase in the fields of genomics, modern biology, human health, and STEM; and the expectation is that they will be better prepared to pursue career opportunities in genomics and/or STEM related fields. Moreover, there is a demand to facilitate a potentially transformative shift in in-school teaching practices that favors a more interdisciplinary and integrational approach to life science instruction [21].

Since many schools throughout the nation, especially in New York City, do not focus on genomics or genomic careers and/or STEM career services, this research will provide a personal connection for underrepresented, urban secondary school students to engage with, and develop a better understanding of, genomic principles and practices in a way that will more fully acclimate them toward these fields and encourage them to pursue the education necessary to participate in those careers. An essential question that was addressed in this research was: What is a progressive method to teach genomics while enhancing a better understanding of biology for secondary school students?

Given the novelty of this course, there was limited prior published research or information on how learning the principles and practices of genomics impacts secondary school students learning, whether its use in a STEM course enhances education. Thus, much of the theory and curriculum design was guided by more general learning theory and best classroom practices combined with judicious selection of genomic principles and practices that most likely would address secondary school students’ interest, given the best evidence to date. Therefore, I used a survey instrument that was administered before and after the course to examine relationships between the use of Likert scale items and student knowledge and attitudes pertaining to genomics. Based on previous research of genomics education in secondary school settings [13, 22–24] this research aims to address the needs of secondary school biology students to make their biological education better grounded in, and informed by, modern advances in genomics; including relationships to human health principles and practices and career awareness in genomics and STEM. The goal is to document and analyze how the participants’ engaged with and learned genomic principles and practices, and how to establish future research that will address the merits of learning genomics; and thus, promote a better understanding of biology, STEM, and human health, more broadly.

## Materials and methods

The research has been approved by the Institutional Review Board (IRB) of Teachers College Columbia University. All of the 25 participants agreed to participate in the course, and the relevant data-gathering methods as part of the course development and documentation in accordance with the rules of the college where the course was held. Teachers College Columbia University approved this previously gathered data as archival evidence to be used in this thesis research (Subject: IRB Approval: 20-169 Protocol Date: 01/10/2020). The students who participated were assigned a number to maintain anonymity, and all information was secured in a locked location to prevent possible dissemination that might lead to identification of the school where the study was done and any of the participants.

Data was gathered by the following means as explained more fully in the following subsections: a) Likert survey of attitudes and critical judgment regarding topics related to genomics, b) Learning outcomes assessment, and c) and interview evidence.

### Participants and research settings

This research was conducted at a public four-year college in New York City, New York. A sample of 25 secondary school students participated in the research, all of which are currently attending various secondary schools throughout New York City, New York. All of the 25 participants were in the age range from 13-15 years (mean = 14, s.d. = 0.93) of age and were enrolled in a free learning opportunity that included this genomic course. It was not evaluated with a grade, nor is it included as part of their formal secondary school required education. Multiple STEM courses are offered including chemistry, anatomy and physiology, and advanced mathematics. The demographics collected from ninth- and tenth-grade participants included gender and age.

### Description of the genomics course: Introduction to genomics

The objective of the course was to develop and design a curriculum that utilizes student-centered, computer-based learning, and other resources from informal learning at AMNH while developing the students’ content knowledge of genomics, biology, human health, STEM, and interdisciplinary skills and strategies. This is an appropriate approach for integrating genomics into the majority of general biology curricula [25], because, this is one of the gaps in current biology curricula [15] offered in the majority of secondary school classrooms across the United States. Throughout the course participants had multiple opportunities to independently and collaboratively learn the principles, practices, and scientific evidence-based aspects, of genomic diseases, including working with “big data” using bioinformatics databases.

> The development of novel bioinformatics approaches, we have gained a deeper understanding of the microbiome and its impact on health and disease in disorders of the skin (psoriasis, acne), the gut (Crohn’s, colon cancer, colitis), the vaginal tract, and the oral cavity (caries) [26].

In addition to becoming more fully informed about genomics, the participants will better understand the nature of science, science investigation, and authentic interdisciplinary learning, and become familiar with the ethical, legal, and social implications (ELSI) associated with genomics. Each of the lessons were developed on vetted design principles of: a) They have clear pedagogical objectives; b) They are integrated with lessons taught in the lecture; c) They are designed to integrate the learning of science content with learning about the process of science; and d) They require student reflection and discussion [19].

The Introduction to Genomics course was developed through insights gained from prior experiences teaching genomic and STEM courses, collaboration with the Sackler Institute for Comparative Genomics at AMNH, and professional development at the Jackson Laboratory. Student professional development and informal learning at AMNH has shown to be effective in stimulating secondary school science learning, interests, and career development in genomics and STEM [27, 24]. One of the main goals was to have the participants engage in lifelong learning of genomics education, conceptualize biological processes, collaborate with developing solutions, and acquire new vocabulary terms from the course that they generally would not have learned in their current secondary school science class. Putting these factors into place would highly support the process of lifelong learning of genomics, biological, and human health concepts. To further test the validity as to why this content is practical, meaningful, interdisciplinary, and applicable to areas of their lives the participants were introduced to relevant websites (S1 Fig). These websites further enhanced their conceptions of genomics as well as addressing the key points mentioned throughout the genomics course. Based on previous observations of Internet use and searches among primary and secondary school students and science teachers, it is essential for the participants to become familiar with reputable websites that are recognized throughout the scientific community. Previous research with cyberlearning has demonstrated that educational computer-based technology has improved students develop a rich comprehension of learning genetics [30].

Utilizing a number of CBL pedagogical skills [31] I focused on the essential terms/vocabulary, genomic concepts and processes to represent clear principles instead of rote facts and details by removing certain terms related to cellular organelles with the exception of the nucleus. Some of the cellular organelles were removed because, although it is essential to learn the cellular organelles to better understand molecular biology, it was more important for students to become familiar with scientific concepts that they have not accustomed to learning.

### Pedagogy

The Biological Sciences Curriculum Study (BSCS) 5E model engage, explore, explain, elaborate, and evaluate provided an effective alternative to traditional and/or autocratic pedagogy [28–29], which tends to focus mainly on one type of activity, processing symbolic information [32]. Using the BSCS approach, the participants were encouraged to use their prior knowledge to engage interest in genomics and related subjects and begin to explore, how these topics occur in the natural world. After acquiring an understanding of genomics, they explain the particular phenomenon they learned and elaborate their understanding through new experiences. Lastly, students evaluate their comprehension and apply it to real-life situations.

### Course structure and design

Each class began with an essential question of the day that was intended to elicit a response from the participants and orient them to the topics for the class session. Answers were listed on the board and briefly elaborated on. Examples include, How has DNA technology impacted law enforcement?; Is it ethical for major sport organizations to sequence an athlete’s genome?; What are three examples of a genetic disease? This was followed by a brief lecture pertaining to the topic of discussion. This included listing genomic terminology, elaborating on biological or scientific processes, or links to society and real-world applications.

To ensure that the course, which was the focus of the research, was best designed for the participants, I used prior genomic resources, curriculum, lesson, pedagogical skills and strategies, and student feedback from a similar genomic course that was taught on three separate occasions. The major difference between the current research and the previous courses taught was, the use of a Likert-scale and in-depth student interviews. The genomics course was taught three times previously before this study was undertaken. However, at that time, none of the relevant evidence from the participants responses to the research instruments was gathered. This prior experience was valuable in designing the course and methodology used in this research study.

During the first session of the course (Day 1) a general questionnaire was administered consisting of background information and prior experience in learning about topics pertaining to genomics and genetics. In addition, a Likert pre-survey was administered to document the participants percepts of ethical, legal, and social topics related to genomics, biology, and human health; and beliefs and attitudes associated with genomics in general and more specifically about its use in scientific research (S2 Fig). Day 1 also focused on measuring the initial status of the students’ knowledge by administering a 25-multiple-choice content achievement test. (S3 Fig), it was constructed to measure the level of prior content knowledge in genomics, biology, and human health concepts. It also provided some background information on the students’ prior knowledge [33] to help guide the more student-centered approach used in the learning experiences. Data gathered from the three replications of the course were compiled into one composite set of results and used to address the research questions posed in this study.

### Likert survey evidence

A Likert survey with 15 items, and five options per item (S2 Fig), was designed to tap some of the critical judgmental, affective, and emotive orientations of the students who participated in the course. The survey consisted of three sections designed to assess the following dimensions: a) opinions about learning modern genomic and genetic principles, b) ethical choices associated with genomics, and c) beliefs and attitudes associated with genomics. The differences in the post-survey responses compared to those in the pre-survey responses provided evidence of changes in these ‘affective dimensions.’

### Learning outcomes assessment

The pre-and-post content achievement test (S3 Fig) consisted of 25 multiple-choice questions of the following topics: Five of the questions (Questions 1-5) addressed basic concepts of genetics [cellular components and processes, chromosome structure and function, inheritance], twenty of the questions (Questions 6-25) addressed genomic principles and practices (analyzing DNA, bioinformatics, HGP, Human Microbiome (HMB), and ELSI). The purpose of administering the pre-test was to gather evidence of the participants’ prior content knowledge pertaining to modern biological, genomics, human health, and ELSI principles as a baseline for comparison with any achievement gains as assessed by the post-test; and to help set the breadth and depth of content that was introduced during the duration of the course. The difference in the means of the pre-test and post-test scores was used to assess student achievement in the course.

### Interview procedures and evidence

Interviews were collected to obtain qualitative evidence beyond that of the Likert surveys. Nine students volunteered to be selected to be respondents; five females and four males. The individual interview with each respondent was held at the very end of the study. The items in the Likert survey served, partially, as a guide to more fully probe students’ perceptions of the course experiences. The questions used in the interview are presented in the (Analysis of participants interviews). Conducting an interview with participants provided additional evidence of the participants’ percepts in the form of expanded (typically qualitative) narrative beyond the qualitative and semi-quantitative data. This was largely intended to provide more open-ended evidence of participants’ reporting of the learning experience and it provided an opportunity for the participants to elaborate on responses made previously to the Likert-scale survey items. Thus, the Likert-scale survey items served as a scaffolding method for the interview questions presented initially to the respondent, as a way of focusing the interview and providing a context to encourage more elaborate narrative by the respondent. The interviews were individually administered, and not audio-taped. Detailed notes were taken including verbatim quotations of the student’s narrative, where appropriate. The respondents’ narrative evidence was used to document and analyze themes that emerged based on my analytical and critical reading of each respondent’s narrative.

### Data analysis and statistical methods

The results of the Likert survey responses, within each if the three sections of the survey, were tabulated as frequencies for each item, and presented as a table for the pre- and post-survey results. Additionally, bar graphs were constructed using Excel as visual evidence of the change in the respondents’ responses toward a more favorable (‘Agree or Completely Agree’) position for each item. This directionality was used as a concise way of exhibiting trends in the respondent’s change in position relative to each Likert item, in addition to the more detailed evidence in the frequency tables. The scores of each student on the multiple response pre- and post-test were tabulated and the means ± standard errors (s. e.) for the pre- and post-test were reported. A paired t-test was used to assess the significance of the mean difference, because the two sets of data are not independent. A p of ≤ 0.05 was used to judge the significance of the t-test results. Evidence from the interviews was qualitative and no statistical analysis was done.

## Results

### Likert survey of attitudes and critical judgment regarding topics related to genomics

The survey consisted of three sections designed to assess the following dimensions: a) opinions about learning modern genomic and genetic principles, b) ethical choices associated with genomics, and c) beliefs and attitudes associated with genomics. The results for the first research question regarding the participants’ opinions about learning modern genomic and genetic principles, ethics, and beliefs and attitudes are presented in Tables 1 to 3 and Figures 1 to 3. Table 1 and Figure 1 present the pre-and post-survey results of the respondents’ opinions for the five items in Section 1 of the survey on the topic of ‘Learning modern genomic and genetic principles.’ Results for Section 1-3 of Likert survey items are listed in the Table 1, 2, and 3; and in Figures 1, 2, and 3.

**Fig 1.**
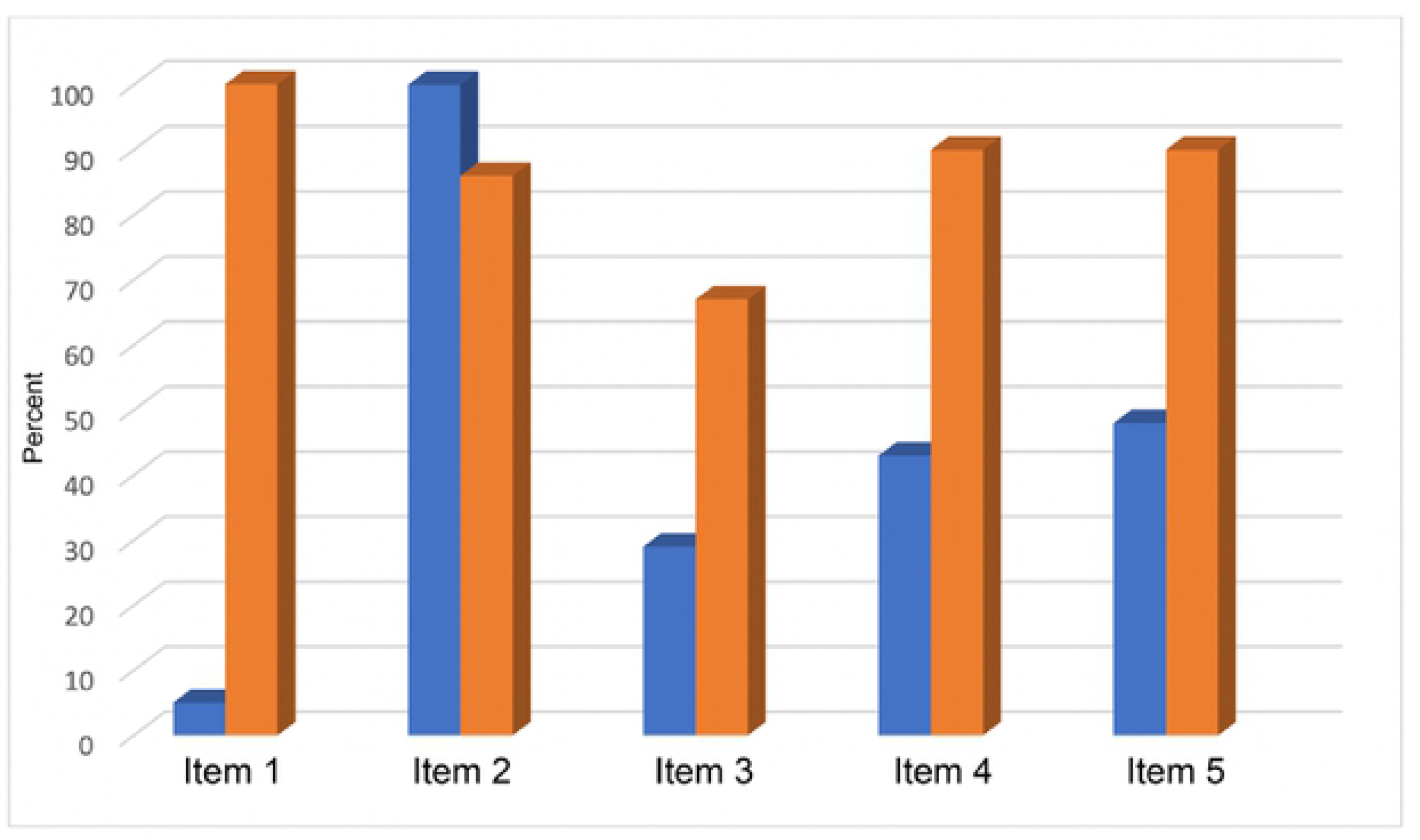
Bar graph of the Likert scale responses for Section 1 Items 1-5. The percentages are shown for the total ‘Agree and Completely agree’ (A and CA) responses for the Likert scale survey (Section 1) items 1-5 (Section 1) related to opinions about learning modern genomic and genetic principles. Pre-survey (blue) and post-survey (red) results.

**Fig 2.**
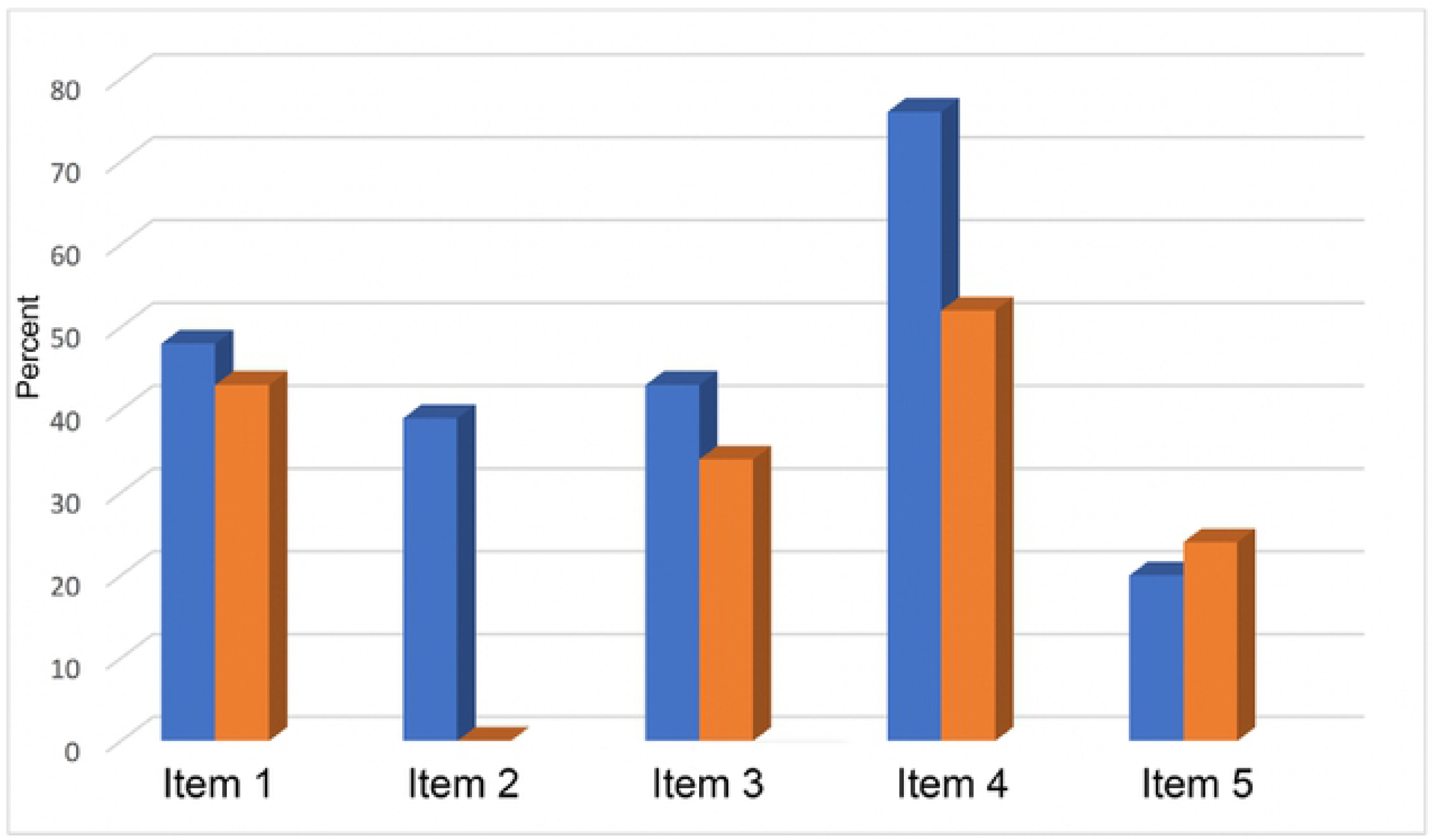
Bar graph of the Likert scale responses for Section 2 Items 1-5. Bar graph showing the percentages of the total ‘Agree’ and Completely agree’ (A and CA) responses for the Likert scale survey (Section 2). Items 1-5 relate to ethical choices associated with genomics. Pre-survey (blue) and post-survey (red) results.

**Fig 3.**
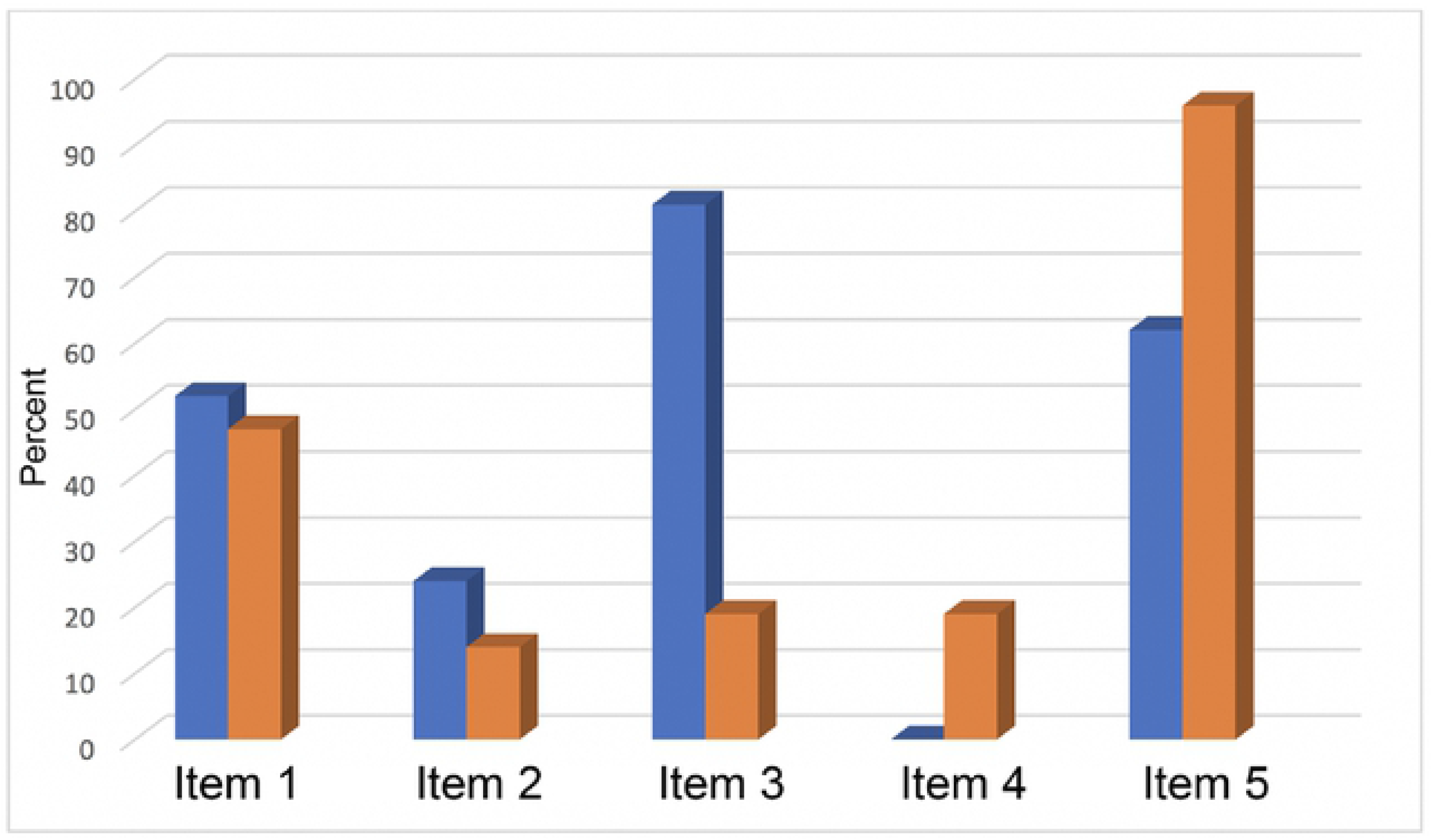
Bar graph of the Likert scale responses for Section 3 Items 1-5. Bar graph showing the percentages of the total ‘Agree and Completely Agree’ (A and CA) responses for the Likert scale survey (Section 3) Items 1-5 related to ‘beliefs and attitudes associated with genomics research and applications.’ Pre-survey (blue) and post-survey [red] results.

**Table 1.**
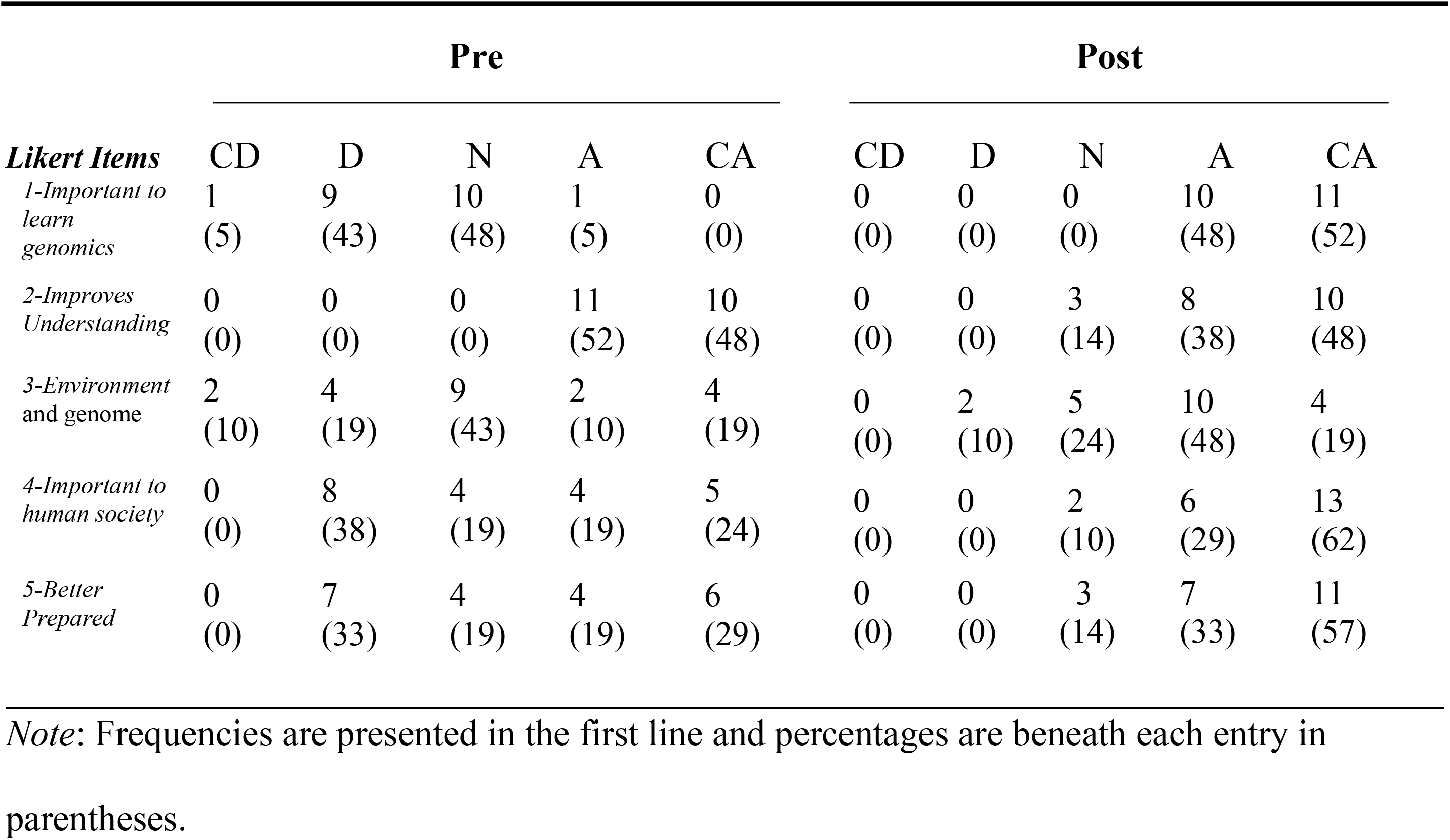
Pre-and-post-Likert survey (Section 1) results of the respondents’ opinions about learning modern genomic and genetic principles.

| *Likert Items* | Pre |  |  |  |  | Post |  |  |  |  |
| --- | --- | --- | --- | --- | --- | --- | --- | --- | --- | --- |
|  | CD | D | N | A | CA | CD | D | N | A | CA |
| *1-Important to learn genomics* | 1<br>(5) | 9<br>(43) | 10<br>(48) | 1<br>(5) | 0<br>(0) | 0<br>(0) | 0<br>(0) | 0<br>(0) | 10<br>(48) | 11<br>(52) |
| *2-Improves Understanding* | 0<br>(0) | 0<br>(0) | 0<br>(0) | 11<br>(52) | 10<br>(48) | 0<br>(0) | 0<br>(0) | 3<br>(14) | 8<br>(38) | 10<br>(48) |
| *3-Environment and genome* | 2<br>(10) | 4<br>(19) | 9<br>(43) | 2<br>(10) | 4<br>(19) | 0<br>(0) | 2<br>(10) | 5<br>(24) | 10<br>(48) | 4<br>(19) |
| *4-Important to human society* | 0<br>(0) | 8<br>(38) | 4<br>(19) | 4<br>(19) | 5<br>(24) | 0<br>(0) | 0<br>(0) | 2<br>(10) | 6<br>(29) | 13<br>(62) |
| *5-Better Prepared* | 0<br>(0) | 7<br>(33) | 4<br>(19) | 4<br>(19) | 6<br>(29) | 0<br>(0) | 0<br>(0) | 3<br>(14) | 7<br>(33) | 11<br>(57) |

*Note:* Frequencies are presented in the first line and percentages are beneath each entry in parentheses.

**Table 2.** Pre-and-post-Likert survey (Section 2) results of the respondents’ ethical choices associated with genomics.

| Likert Items | Pre |  |  |  |  | Post |  |  |  |  |
| --- | --- | --- | --- | --- | --- | --- | --- | --- | --- | --- |
|  | CD | D | N | A | CA | CD | D | N | A | CA |
| 1-Correct<br>"Bad genes" | 1<br>(5) | 3<br>(14) | 7<br>(33) | 5<br>(24) | 5<br>(24) | 2<br>(10) | 5<br>(24) | 5<br>(24) | 6<br>(29) | 3<br>(14) |
| 2-Alter the<br>genome | 3<br>(14) | 1<br>(5) | 9<br>(43) | 4<br>(19) | 4<br>(19) | 6<br>(29) | 12<br>(57) | 3<br>(14) | 0<br>(0) | 0<br>(0) |
| 3-Increase<br>life span | 8<br>(38) | 0<br>(0) | 4<br>(19) | 6<br>(29) | 3<br>(14) | 3<br>(14) | 5<br>(24) | 6<br>(29) | 6<br>(29) | 1<br>(5) |
| 4-Baby's gene | 2<br>(10) | 0<br>(0) | 3<br>(14) | 9<br>(43) | 7<br>(33) | 0<br>(0) | 4<br>(19) | 6<br>(29) | 8<br>(38) | 3<br>(14) |
| 5-Neanderthal | 8<br>(38) | 6<br>(29) | 3<br>(14) | 2<br>(10) | 2<br>(10) | 5<br>(24) | 6<br>(29) | 5<br>(24) | 3<br>(14) | 2<br>(10) |

*Note:* Frequencies are presented in the first line and percentages are beneath each entry in parentheses.

**Table 3.** Pre-and-post-Likert survey [Section 3] results of the respondents’ beliefs and attitudes associated with genomics.

|  | Pre |  |  |  |  | Post |  |  |  |  |
| --- | --- | --- | --- | --- | --- | --- | --- | --- | --- | --- |
| *Likert Items* | CD | D | N | A | CA | CD | D | N | A | CA |
| *1-Medical Research* | 0<br>(0) | 2<br>(10) | 8<br>(38) | 4<br>(19) | 7<br>(33) | 2<br>(10) | 4<br>(19) | 5<br>(24) | 7<br>(33) | 3<br>(14) |
| *2-Genetic research* | 3<br>(14) | 5<br>(24) | 8<br>(38) | 5<br>(24) | 0<br>(0) | 9<br>(43) | 7<br>(33) | 2<br>(10) | 3<br>(14) | 0<br>(0) |
| *3-Microscopic* | 0<br>(0) | 2<br>(10) | 2<br>(10) | 9<br>(43) | 8<br>(38) | 7<br>(33) | 5<br>(24) | 5<br>(24) | 4<br>(19) | 0<br>(0) |
| *4-DNA database* | 9<br>(43) | 9<br>(43) | 3<br>(14) | 0<br>(0) | 0<br>(0) | 5<br>(24) | 7<br>(33) | 5<br>(24) | 4<br>(19) | 0<br>(0) |
| *5-Polygenetic diseases* | 3<br>(14) | 2<br>(10) | 3<br>(14) | 4<br>(19) | 9<br>(43) | 0<br>(0) | 0<br>(0) | 1<br>(5) | 6<br>(29) | 14<br>(67) |

*Note:* Frequencies are presented in the first line and percentages are beneath each entry in parentheses.

### Results for Section 1 of Likert survey

The pre and postsurvey results for the responses to items in Section 1 of the Likert survey related to learning modern genomic and genetic principles are presented in Table 1.1 and Figure 1.1.

### Results for Section 2 of Likert Survey

The pre- and post-survey results for the responses to items in Section 2 of the Likert survey related to ethical choices associated with genomics are presented in Table 2 and Figure 2.

Figure 2 provides visual evidence of the changes in the respondents’ percepts from the Likert pre-to post-survey for items 1 to 5 of Section 2, particularly illustrating the wide variability in the magnitude of the response change, while illustrating that most of the changes were more negative (Items 1 to 4).

### Results for Section 3 of Likert survey

Section 3 of the Likert survey focused more specifically on the use of genomics in research, including aspects of research data, possible medical remedial procedures, genomic diseases, and use of comprehensive data bases containing individual DNA evidence. The results of the Likert pre- and post-surveys are presented in Table 3 and Figure 3. Items 1 to 3 show varying degrees of decrease in positive percepts; while Items 4 and 5 show varied increases in positive orientation toward strongly ‘Agree.’

The data in Table 3 is presented visually in Figure 3, particularly focusing on the change from, pre-to post-survey for Items 1 to 5 of Section 3 in the Likert survey. The general trend toward a more favorable response to Item 5, and a decline in positive perspective for Item 3, is particularly evident.

### Analysis of composite results for sections 1 to 3 of Likert survey

Overall, there was a net increase in the percentages tending toward a positive response to Items 1 to 5 in Section 1 (1.1 to 1.5, Figure 4) on ‘opinions about learning modern genomic and genetic principles’ (mean ± s.e. = 38.6 ± 17.2). For Items 1 to 5 in Section 2 (2.1 to 2.5, Figure 4) on ‘ethical choices associated with genomics,’ the net change was toward more negative (-14.4 ± 7.3). The response to items 1 to 5 in Section 3 (3.1 to 3.5, Figure 4) was mixed, with a small net negative change in percent toward less approval to the items (−4.8 ± 16.4). The relatively large s.e. indicates the diversity in the responses, particularly with items in Sections 2 and 3 being more negative than those in Section 1.

**Fig 4.**
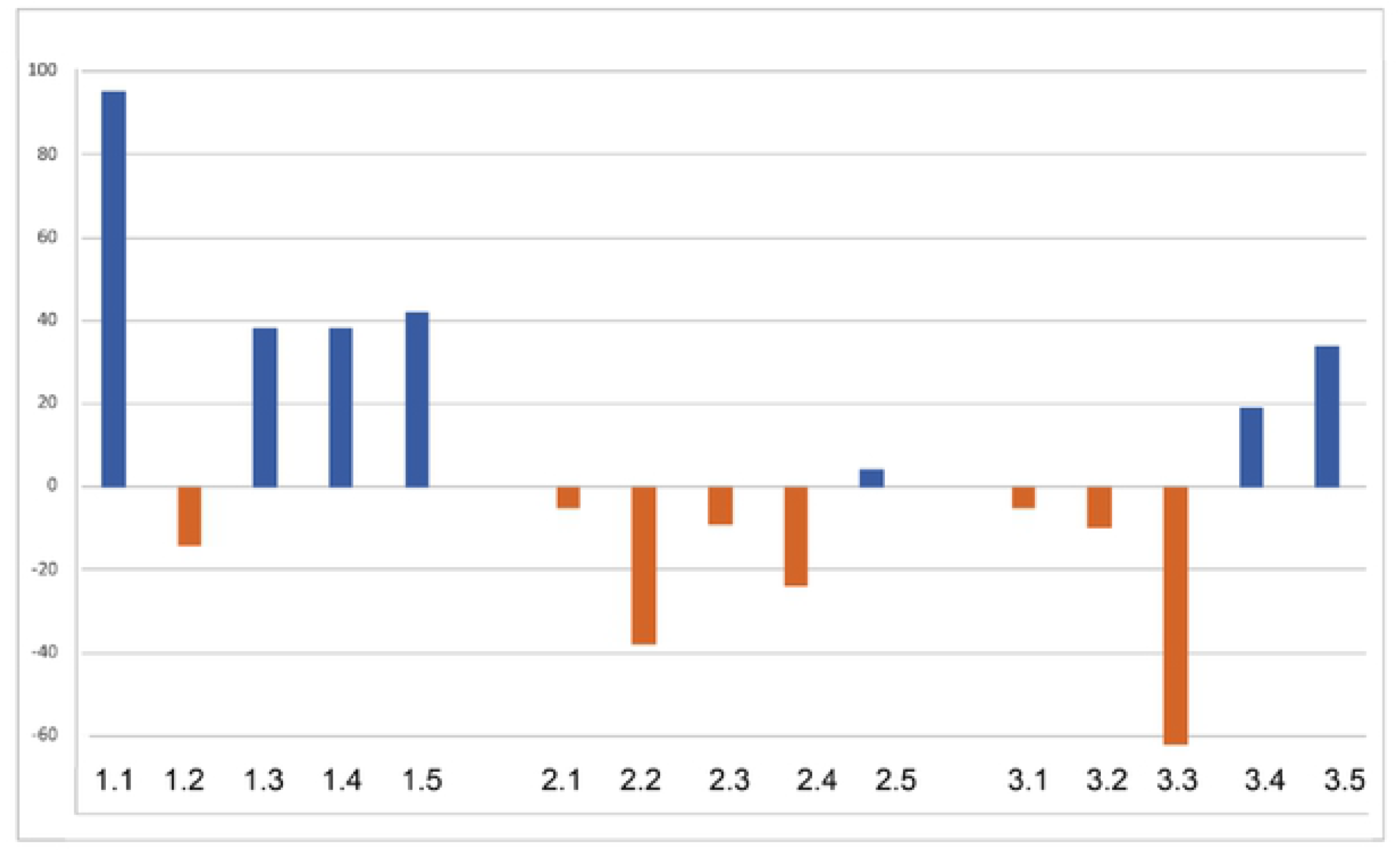
Net percent change toward more positive responses (blue bars) or toward less positive responses (red bars) to the Likert survey items in Sections 1 to 3. Section 1 (1.1 to 1.5) on ‘opinions about learning modern genomic and genetics principles,’; Section 2 (2.1 to 2.5) on ‘ethical choices associated with genomics; and Section 3 (3.1 to 3.5) on ‘beliefs and attitudes associated with genomics research and applications.’

### Analysis of learning outcomes assessment

The results for the second research question addressed the following: The content achievement gains reported in this study, for the difference in pre-test and post-test scores, indicate that a student-centered approach with CBL can be a productive way to increase student understanding of broad-based issues in genomics.

The mean scores ± standard errors are 61.25 ± 1.19 for the pre-test and 83.75 ± 1.03 for the post-test. The paired *t*-test for the difference in the means between the pre-and post-test was highly significant, t =19.3 (p < 0.0001, df =23). Overall, the mean gain was 22 scale points within a 100-point total score.

### Analysis of participants interviews

Results for the third research question addressed evidence related to the participants’ perceptions of their learning experience. There were five interview questions, and nine participants who were interviewed.

#### Interview question 1

What were some interesting topics that you learned throughout the duration of the course? Of the nine participants that were interviewed about their interests in the course, all of them did find the course to be interesting and the course did clarify some genetic concepts that they previously misunderstood. They were also asked if there were any genomic terms that were unfamiliar to them prior to the course, and they all answered yes. The responses that came up most frequently were: Genome-Wide Association Studies (GWAS), ELSI, Genetic Information Nondiscrimination Act (GINA), micropipetting, HMB, pharmacogenomics, and epigenetics.

#### Interview question 2

Throughout this course we learned and focused on polygenetic and monogenetic diseases. Was this a topic that you found interesting? All of the nine participants agreed that learning this topic helped clarify and dispel some misconceptions pertaining to inherited and acquired diseases. Including categories such as: sex-linked, autosomal dominant, autosomal recessive, and chromosomal abnormalities. Moreover, there were specific diseases that were of particular interest, which optimized their proclivity to enhance knowledge.

#### Interview question 3

Did you enjoy utilizing CBL throughout the course? Given the innovative and interactive nature of CBL, all of the nine participants were very comfortable and stated that their learning was influenced due to the student-centered learning framework (learning at their own pace as an autonomous learner).

#### Interview question 4

Were there any particular topics that you enjoyed learning regarding ELSI? Each of the interviewee responses demonstrated that they were aware of and believed that they were very successful in making intuitive decisions towards ELSI of genomics. Many of the participants had varying interests towards a particular topic of ELSI, including: genetic discrimination, designer babies, personal genomics and healthcare, and genetic engineering. One specific example was with Item 4 on Section 3 of the Likert survey that addressed the issue of maintaining public data bases on human DNA information (Table 4.3). Seven of the nine participants interviewed ‘Completely disagreed’ with Item 4. An excerpt during the interview with participant #12 highlights the lack of ELSI at their current school. The following are excerpts from the interview responses that pertain to Interview question 4; the prefix “R” represents the interviewer’s comments or questions, and each respondent’s answer is coded with the respondent’s participant number (#).

R: You mentioned that you were interested in ELSI, any particular topic that you were interested in learning?
#12: Yea I really liked when we did our presentations, I did mine on the DNA database. I think most of us enjoyed picking our own topic, but when you showed us about the Golden State Killer, it really was interesting to learn how the world is changing.
R: What do you mean by that?
#12:Like how the government and society is being watched and there really isn’t as much privacy as people think. I just think it was cool learning about this more often. At first, I was total against it, but now I think there is some proof to why it’s being done in the UK. I pretty sure that from what we looked at on the websites that we are all going to have our info in some database. Like now that you told us about the inheritance websites, I told a few of my friends that it is not worth doing.
R: OK, why?
Because we are all going to be watched or someone will be able to hack us or something like that.
R: Do you learn about ELSI in your school?
#12: No, I wish we did. I like talking and learning about ethics and how it affects society. We do a lot of labs and cool activities, but we don’t use the websites that you showed us or get to talk about many things that we don’t realize. Like when we were talking about gene editing and the designer babies, it was just cool to know that this type of science exists.

#### Interview question 5

There were two days that we focused on genomic careers. Do you think that you would pursue a career in genomics? Seven of the nine participants explained that they may continue to enroll in similar STEM courses that focus on student-centered learning and genomic/STEM careers. They realized that they were ultimately responsible for their career trajectory, however the proper resources were not available at their school or their science teachers or guidance counselors were novices in STEM/education and college placement for STEM careers. Several expressed they were inspired and would study genomics in college.

The following are excerpts from the interview responses that pertain to Interview question 5. (R=researcher):

R: Do you believe that this course prepared you to enroll in a career in genomics?
#20: Oh yes definitely. It’s a very interesting subject.
R: Can you elaborate?
#20: Well all of the topics that we learned are interesting and I think that I have what it takes to become a scientist. I really liked when you learned about the different careers from the website. Most of those careers I didn’t know existed and we definitely don’t learn them in our other science class.
R: Which genomic career are you interested in pursuing?
#20: Probably forensics or something involved in law and government. I mean what we learned about ELSI was also very interesting so I think that would be another career choice. You see that is the dilemma that I have sometimes and I have no one at my school to help. They are good teachers and guidance counselors but there are so many students.

## Discussion

The participants in this study were 25 volunteer secondary school students from public schools in New York City. All 25 students completed the genomics pre- and post-achievement test, 21 completed the pre- and post-Likert surveys, and nine were interviewed to gain insight into their perceptions of the content in the course. The results for each of the three research questions will be discussed in sequential order, including a cross comparative analysis and discussion of the results for the three Likert-survey items that were used in addressing Research Question 1. Finally, the last sections include: Implications and Relationships to the Literature, Summary of Strengths and Limitations, Including Transferability and Scalability, Implications for Future Research and Applications, and conclusions.

### Implications and relationships to the literature

New approaches for teaching genomic principles and practices acquired during the learning process is critical in the hopes of producing scientifically literate citizens. This research demonstrates that understanding the principles and practices of genomics by using a student-centered approach with limited lecturing, a CBL, and interdisciplinary learning, and assessment tools aimed at promoting interest in genomics can substantially influence students’ critical thinking about, and interest in, important societal and scientific issues in modern genomics. The success of the Introduction to Genomics course can be attributed to the implementation of sufficient structured experiences in addition to more student-centered learning, such as being explicit to the participants about which skills and strategies are specifically important for progress in STEM education, introducing and utilizing scientific terminology, and encouraging student collaboration. These skills and strategies also place an emphasis on STEM careers and scientific literacy [34].

Overall the analysis of the participants responses to the Likert scale, pre-and-post-tests assessments, and interviews revealed that they expressed an increasing interest in genomic and STEM education, especially in ELSI awareness. Among the topics that the participants believed were of interest included: GINA, direct-to-consumer tests, DNA databases, reproductive issues, and newborn screening inspired them to pursue the education necessary to acclimate themselves in a scientific career. Additionally, the logical alignment of the curriculum, pedagogy, and assessment procedures that were used support the validity and reliability of the research [35]. Moreover, it permitted the participants to collaborate with one another in an academic and urban setting that of which contributed to learning STEM.

Notwithstanding the rather substantial achievement gains [pre-and-post-test] and increased interest in genomics (Likert survey), some of the participants during the interview were highly responsive to discuss in what ways they were deficient in their knowledge of genomics as well as the areas where they were more proficient, as is consistent with findings from other prior research [35]. Most of the participants did not have access to educational resources for learning genomics, and were not being exposed to these topics in their current schooling. Yet they demonstrated in this study, the proclivity necessary to acquire the content knowledge effectively. For example, Participant #12 attends an elite NYC public secondary school, notably one of the most prominent and selective in the county that specializes in STEM education and has a double period, college-level, semester long genetics course provided for 12^th^ graders. The majority of the course is laboratory-based utilizing modern scientific applications including micropipetting techniques, bacterial transformation, electrophoresis, PCR, chromatography, and DNA extraction. While the course is progressive with modern principles, it is like many other science courses that exclude interdisciplinary topics such as ELSI. These interdisciplinary connections made possible with properly organized genomics lessons can connect students to the pragmatics of the real-world, and encourages them to voice their opinions and develop critical thinking skills [3,14–15].

### Summary of strengths and limitations

The development of the Introduction to Genomics course and curriculum was designed for the duration of one year to allow secondary school students to acquire content knowledge and familiarity with modern biological principles and practices, emphasizing genomics, in a STEM college setting. Overall, the participants enjoyed CBL as it encouraged group discussions and presentations, which helped in retaining information and improving interdisciplinary and cognitive skills. Student-centered pedagogy was generally very positive. The majority of participants stated that they preferred the BSCS 5E model as opposed to traditional/autocratic pedagogy that was typical in their schools. The report Vision and Change in Undergraduate Biology Education: A Call to Action American Association for the Advancement of Science [36] expressed great concern for an increase in more student-centered learning in undergraduate biology and science education classes. Many of the ELSI components associated with genomics were, at first, unfamiliar or seemed esoteric to them; however, exposure to these topics was very transformative, especially since many had differing critical judgements before and after the course.

There were many informal components that were implemented into the classroom, including micropipetting, DNA extraction, and AMNH laboratory techniques via weblinks. However, there were no visits to any informal learning setting such as AMNH or the Dolan DNA Center (Cold Spring Harbor Laboratories). Informal learning provides opportunities to engage students with authentic and layered learning [all ages learn different levels of content] experiences, and connect students to real-world applications. For the student, informal learning exposes them with rigorous learning opportunities beyond what they would learn in their classroom, and allows them to potentially interact with curators, researchers, and staff members [37]. While CBL was implemented on a daily basis, there was absence of a wet lab that is also useful for conceptualizing genomic content. The course itself was not conducive to employing hands-on laboratory assignments, being that it was in a typical college classroom. Further genomics curricula, based on innovative approaches such as the one introduced here, may be improved by including appropriately selected laboratory experiences to provide a deeper understanding of the practices of science associated with genomics research and applications.

As a science education researcher, it is important to identify what students already know about genomics, how they represent what they know cognitively, and what experiences, if any, they already have. For example, on the first day of the course, as I have done, and continue to do in up-and-coming STEM courses, I invite comments, and ask students “What do you already know about genomics?” or “What prior knowledge do you already have associated with microbiology?” After eliciting responses and documenting them on the board, I briefly relate new information and explain some relationships to their prior knowledge and highlight the content of the syllabus. However, some students may not have any prior knowledge of the relationship between what they already know and what they will know. Therefore, it is important to include content material in the syllabus that demonstrates the merging of their prior knowledge and newly acquired knowledge of the course.

Prior to the course, some of the participants had a lack of understanding of genomic relationships with other biological and scientific disciplines due to their misunderstanding of the genomics composition of the eukaryotic cell. They encountered difficulties distinguishing between polygenetic and monogenetic diseases, beneficial and harmful bacteria, inheritance, and genomic terminology. In fact, the very first question I ask participants before teaching any course related to genomics is “raise your hand if you have heard of the Human Genome Project.” Based on the low number of hands raised, which unfortunately over the last several years has been less than five, there is an indication that a progressive paradigm shift in genomic education at the secondary school level is indeed essential.

### Transferability

This course was developed for a select group of students from inner city schools and all were volunteers. However, this was a suitable group to examine the introduction of a new approach to biological education such as this one. It is often preferable to begin with a more tractable group of participants to initiate a novel educational experience to better establish likely boundary conditions for its transfer to other learning situations. I believe that this course can benefit colleges and secondary schools not only through dissemination of the research and experiences, but also by implementing the course as a unit or as an elective course. The course is available online at this URL: https://drawingandgenomics.wixsite.com/drawingandgenomics: included other interactive websites and resources that can provide biology or health educators resources to increase genomic awareness, literacy, and knowledge.

While the importance of learning genomics is widely acknowledged in science and science education fields [8,15] challenges remain, especially in the ability for science teachers to apply, enhance, and implement genomic content knowledge in their classrooms, which is a major goal in science education [19]. To achieve this goal in secondary school education, schools and science teachers should enrich their students with topics that address their interests through CBL, student-centered learning, informal learning, and project-based learning.

### Scalability

Students learn better by communicating through collaborating, talking, and interacting because each requires high levels of thinking [38]. Teaching and or establishing a genomics course for a class greater than 30, which I have done previously in a similar academic setting, is also productive, and easily conceivable and applicable. Utilizing the pedagogical skills [as previously mentioned] encourages students to become aware of their pacing, academic responsibilities, prompts them to think more actively, obtain more information via CBL or collaboration to clarify their new and existing knowledge [28]. By structuring the class into groups [whether the class size is 20, 25, or 30 students] increases their potential to provide a feeling of inclusion, community, and collaboration for many students who may otherwise feel isolated in biology classrooms [38].

## Conclusion

By engaging secondary school students in a modern genomics course as documented here, they were given the opportunity to develop more concise knowledge and critical thinking skills about a unique STEM domain, while learning how to engage in science that is contemporary and applicable to real-life/real-world issues. Given that the current curriculum in science classes reflects domains in science that are relatively out-of-date, it is important for students to engage in science that reflects cutting-edge discoveries including, personalized medicine and direct-to-consumer; and domains in science that have an application to real-life phenomena. Genomics offers this type of real-life/real-world applications that encourage all students at all academic levels to conceptualize genomic diseases, medicine, ethics, beliefs, research, and careers.

The results of this research showed that genomic curricular materials and resources are, in fact, available and improving to include issues addressing individualization and society. In addition, this research focused on using CBL and interdisciplinary science learning, as well as, diverse and innovative teaching [28,38], and assessment strategies. While the clear benefits of teaching genomics has been well-supported throughout this research, the continuation of integrating aspects of genomics into to the majority of secondary school curricula has yet to be implemented into science classrooms. Since genomics and genetics both have foundational science and real-life/real-world applications, the content is well-grounded to be integrated in an interdisciplinary way with other foundational sciences, especially STEM courses. Moreover, learning genomics encourages students to become aware of modern medical advancements and it is necessary and beneficial for genomics to be incorporated into to secondary schools [3,14–15].

Currently, medical, nursing, pharmacology, and other human health programs are gradually exposing students to a curriculum that introduces them to genomic principles and practices [as previously mentioned]. As science educators, it is our obligation to inform the next generation of future scientists to be knowledgeable in genomics so they can transform new information and discoveries into scientific practice.

## Acknowledgements

The author would like to thank the secondary school participants and the Jackson Laboratory (Bar Harbor, ME) that made this research possible.

## Supporting information

**S1 Fig. Genomic Educational Resources and Interactive Websites**

**S2 Fig. Likert-Scale Survey Items**

## References

1. Collins FS, McKusick VA. Implications of the Human Genome Project for Medical Science. JAMA. 2001;285(5):540–544. doi:10.1001/jama.285.5.540

2. Dougherty MJ. Closing the gap: inverting the genetics curriculum to ensure an informed public. Am J Hum Genet. 2009 Jul;85(1):6–12. doi: 10.1016/j.ajhg.2009.05.010. Epub 2009 Jun 25. PMID: 19559400; PMCID: PMC2706960.

3. Dressler LG, Jones SS, Markey JM, Byerly KW, Roberts MC. Genomics education for the public: perspectives of genomic researchers and ELSI advisors. Genet Test Mol Biomarkers. 2014 Mar;18(3):131–40. doi: 10.1089/gtmb.2013.0366. Epub 2014 Feb 4. PMID: 24495163; PMCID: PMC3948600.

4. Bybee R. What Is STEM Education? Science. 2010; 329(5995), 996.

5. Dougherty MJ, Pleasants C, Solow L, Wong A, Zhang H. A comprehensive analysis of high school genetics standards: are states keeping pace with modern genetics? CBE Life Sci Educ. 2011 Fall;10(3):318–27. doi: 10.1187/cbe.10-09-0122. PMID: 21885828; PMCID: PMC3164571.

6. Knippels, M.C.P., Waarlo, A.J. and Boersma, K.T., 2005. Design criteria for learning and teaching genetics. Journal of Biological Education, 39(3), pp.108–112.

7. Kung JT, Gelbart ME. Getting a head start: the importance of personal genetics education in high schools. Yale J Biol Med. 2012 Mar;85(1):87–92. Epub 2012 Mar 29. PMID: 22461746; PMCID: PMC3313542.

8. Hood L, Rowen L. The Human Genome Project: big science transforms biology and medicine. Genome Med. 2013 Sep 13;5(9):79. doi: 10.1186/gm483. PMID: 24040834; PMCID: PMC4066586.

9. Collins FS. Medical and societal consequences of the Human Genome Project. N Engl J Med. 1999; 341(1), pp.28–37.

10. Cook-Deegan, Robert M. The gene wars: Science, politics, and the human genome. WW Norton & Company. 1994.

11. Collins F, Green E, Guttmacher AE, Guyer MS. A vision for the future of genomics research. Nature. 2003; 422. 835–847. https://doi.org/10.1038/nature01626

12. Venter JC, Cohen D. The Century of Biology. NPQ. 2014 31(1), 28–37.

13. Bigler AM, Hanegan NL. Student content knowledge increases after participation in a hands-on biotechnology intervention. J Sci Educ Tech. 2011; 246–257.

14. LaRue KM, McKernan MP, Bass KM, Wray CG. Teaching the Genome Generation: Bringing Modern Human Genetics into the Classroom Through Teacher Professional Development. J STEM Outreach. 2018 Apr;1(2):48–60. doi: 10.15695/jstem/v1i1.12. Epub 2018 May 3. PMID: 31667467; PMCID: PMC6821449.

15. Wefer SH, Sheppard K. Bioinformatics in high school biology curricula: a study of state science standards. CBE Life Sci Educ. 2008 Spring;7(1):155–62. doi: 10.1187/cbe.07-05-0026. PMID: 18316818; PMCID: PMC2262119.

16. Elmesky R. Building capacity in understanding foundational biology concepts: A K-12 learning progression in genetics informed by research on children’s thinking and learning. Res Sci Educ. 2013;43(3), 1155–1175.

17. Hurle B, Citrin T, Jenkins JF, Kaphingst KA, Lamb N, Roseman JE, et al. What does it mean to be genomically literate?: National Human Genome Research Institute Meeting Report. Genet Med. 2013 Aug;15(8):658–63. doi: 10.1038/gim.2013.14. Epub 2013 Feb 28. PMID: 23448722; PMCID: PMC4115323.

18. Dawson V, Carson K, Venville G. Genetics curriculum materials for the 21st century. Teaching Science. 2010; 56(4), 38–42.

19. Banta LM, Crespi EJ, Nehm RH, Schwarz JA, Singer S, Manduca CA, et al. Integrating genomics research throughout the undergraduate curriculum: a collection of inquiry-based genomics lab modules. CBE Life Sci Educ. 2012 Fall;11(3):203–8. doi: 10.1187/cbe.11-12-0105. PMID: 22949416; PMCID: PMC3433288.

20. Ditty JL, Kvaal CA, Goodner B, Freyermuth SK, Bailey C, Britton RA, et al. Incorporating genomics and bioinformatics across the life sciences curriculum. PLoS Biol. 2010 Aug 10;8(8):e1000448. doi: 10.1371/journal.pbio.1000448. PMID: 20711478; PMCID: PMC2919421.

21. Weber KS, Jensen JL, Johnson SM. Anticipation of Personal Genomics Data Enhances Interest and Learning Environment in Genomics and Molecular Biology Undergraduate Courses. PLoS One. 2015 Aug 4;10(8):e0133486. doi: 10.1371/journal.pone.0133486. PMID: 26241308; PMCID: PMC4524698.

22. Santschi L, Hanner RH, Ratnasingham S, Riconscente M, Imondi R. Barcoding life’s matrix: translating biodiversity genomics into high school settings to enhance life science education. PLoS Biol. 2013;11(1):e1001471. doi: 10.1371/journal.pbio.1001471. Epub 2013 Jan 29. PMID: 23382648; PMCID: PMC3558426.

23. Corn J, Pittendrigh BR, Orvis KS. Genomics Analogy Model for Educators (GAME): from jumping genes to alternative splicing. J Biol Educ (Society of Biology). 2004; 39(1), 24–26.

24. Munn M, Skinner PON, Conn L, Horsma HG, Gregory P. The involvement of genome researchers in high school science education. Genome Research. 1999; 9(7), 597–607.

25. Machluf Y, Yarden A. Integrating bioinformatics into senior high school: design principles and implications. Briefings In Bioinformatics. 2013;14(5), 648–660.

26. Waldor MK, Tyson G, Borenstein E, Ochman H, Moeller A, Finlay BB, et al. Where next for microbiome research? PLoS Biol. 2015 Jan 20;13(1):e1002050. doi: 10.1371/journal.pbio.1002050. PMID: 25602283; PMCID: PMC4300079.

27. Cvijovic M, Höfer T, Aćimović J, Alberghina L, Almaas E, Besozzi D, et al. Strategies for structuring interdisciplinary education in Systems Biology: an European perspective. NPJ Syst Biol Appl. 2016 May 26;2:16011. doi: 10.1038/npjsba.2016.11. PMID: 28725471; PMCID: PMC5516850.

28. Bybee RW, Van Scotter P. Reinventing the Science Curriculum. Educational Leadership. 2006; 64(4), 43–47.

29. Rodriguez S, Allen K, Harron J, Qadri SA. Making and the 5E Learning Cycle. Science Teacher. 2019; 86(5), 48–55.

30. Southworth M, Mokros J, Dorsey C, Smith R. The case for cyberlearning: genomics (and dragons) in the high school biology classroom. The Science Teacher. 2010; 77(7), 28.

31. Clarke, Alan. Designing computer-based learning materials. Gower Publishing, Ltd., 2001.

32. Collins J. Lifelong learning in the 21st century and beyond. Radiographics. 2009; 613–622.

33. Latimier A, Riegert A, Peyre H, Ly ST, Casati R, Ramus F. Does pre-testing promote better retention than post-testing?. NPJ science of learning. 2019; 4(1), pp.1–7.

34. United States. Department of Education. Office of Educational Technology. Transforming American education: Learning powered by technology. 2010.

35. Prozesky D. Assessment of learning. Community Eye Health. 2001;14(38):27–8. PMID: 17491913; PMCID: PMC1705919.

36. American Association for the Advancement of Science. Vision and change in undergraduate biology education: A Call to Action. 2011

37. McQueen J, Wright JJ, Fox JA. Design and implementation of a genomics field trip program aimed at secondary school students. PLoS Comput Biol. 2012;8(8):e1002636. doi: 10.1371/journal.pcbi.1002636. Epub 2012 Aug 30. PMID: 22956895; PMCID: PMC3431290.

38. Tanner KD. Structure matters: twenty-one teaching strategies to promote student engagement and cultivate classroom equity. CBE Life Sci Educ. 2013 Fall;12(3):322–31. doi: 10.1187/cbe.13-06-0115. PMID: 24006379; PMCID: PMC3762997.

